# Division of labour between different PP2A-B56 complexes during mitosis

**DOI:** 10.1101/425579

**Authors:** Giulia Vallardi, Lindsey A Allan, Lisa Crozier, Adrian T Saurin

## Abstract

PP2A-B56 is a serine/threonine phosphatase complex that regulates several major mitotic processes, including sister chromatid cohesion, kinetochore-microtubule attachment and the spindle assembly checkpoint. We show here that these key functions are divided between B56 isoforms that localise differentially to either the centromere or kinetochore. The centromeric B56 isoforms rely on a specific interaction with Sgo2, whereas the kinetochore isoforms bind preferentially to BubR1 and other proteins containing an LxxIxE motif. In addition to these selective interactions, Sgo1 also contributes to both localisations by collaborating with BubR1 to maintain B56 isoforms at the kinetochore and helping to anchor the Sgo2/B56 complex at the centromere. A series of chimaeras were used to map the critical region in B56 to a small C-terminal loop that specifies which interactions are favoured and therefore defines where B56 isoforms localise during prometaphase. Together, this study describes how different PP2A-B56 complexes utilise isoform-specific interactions to control distinct processes during mitosis.

## INTRODUCTION

Protein Phosphatase 2A (PP2A) is a major class of serine/threonine phosphatase that is composed of a catalytic (C), scaffold (A) and regulatory (B) subunit. Substrate specificity is mediated by the regulatory B subunits, which can be subdivided into four structurally distinct families: B (B55), B’ (B56), B” (PR72) and B”’ (Striatin) (Seshacharyulu et al., 2013).

In humans, the B subunits are encoded by a total of 15 separate genes which give rise to at least 26 different transcripts and splice variants; therefore, each of the four B subfamilies are composed of multiple different isoforms (Seshacharyulu et al., 2013). Although these isoforms are thought to have evolved to enhance PP2A specificity, there is still no direct evidence that isoforms of the same subfamily can regulate specific pathways or processes. Perhaps the best indirect evidence that they can comes from the observation that B56 isoforms localise differently during mitosis (Bastos et al., 2014; Nijenhuis et al., 2014). However, even in these cases, it is still unclear how this differential localisation is achieved or why it is needed.

We addressed this problem by focussing on prometaphase, a stage in mitosis when PP2A activity is essential to regulate sister chromatid cohesion (Kitajima et al., 2006; Riedel et al., 2006; Tang et al., 2006), kinetochore-microtubule attachments (Foley et al., 2011; Kruse et al., 2013; Suijkerbuijk et al., 2012; Xu et al., 2013) and the spindle assembly checkpoint (Espert et al., 2014; Nijenhuis et al., 2014). Crucially, all of these mitotic functions are controlled by PP2A-B56 complexes that localise to either the centromere or the kinetochore.

The kinetochore is a multiprotein complex that assembles on centromeres to allow their physical attachment to microtubules. This attachment process is stochastic and error-prone, and therefore it is safeguarded by two key regulatory processes: the spindle assembly checkpoint (SAC) and kinetochore-microtubule error-correction. The SAC preserves the mitotic state until all kinetochores have been correctly attached to microtubules, whereas the error-correction machinery removes any faulty microtubule attachments that may form (Saurin, 2018). The kinase Aurora B is critical for both processes because it phosphorylates the kinetochore-microtubule interface to destabilise incorrectly attached microtubules and it reinforces the SAC, in part by antagonising Knl1-PP1, a kinetochore phosphatase complex needed for SAC silencing (Saurin, 2018). These two principal functions of Aurora B are antagonised by PP2A-B56, which localises to the Knl1 complex at the outer kinetochore by binding directly to BubR1 (Foley et al., 2011; Kruse et al., 2013; Suijkerbuijk et al., 2012; Xu et al., 2013). This interaction is mediated by the B56 subunit, which interacts with a phosphorylated LxxIxE motif within the kinetochore attachment regulatory domain (KARD) of BubR1 (Wang et al., 2016a; Wang et al., 2016b).

As well as localising to the outer kinetochore, PP2A-B56 also localises to the centromere by binding to shugoshin 1 and 2 (Sgo1/Sgo2) (Kitajima et al., 2006; Riedel et al., 2006; Rivera et al., 2012; Tang et al., 2006; Tanno et al., 2010; Xu et al., 2009). The crystal structure of Sgo1 bound to PP2A-B56 has been solved to reveal a bipartite interaction between Sgo1 and the regulatory and catalytic subunits of the PP2A-B56 complex (Xu et al., 2009). This interaction is thought to allow centromere-localised PP2A-B56 to counteract various kinases, such as Aurora B, which remove cohesion rings from chromosome arms during early mitosis in higher eukaryotes (Marston, 2015). The result is that cohesin is specifically preserved at the centromere where it is needed to resist the pulling forces exerted by microtubules. As well as preserving cohesion at the centromere, PP2A-B56 is also thought to balance the net level of Aurora B activation in this region (Meppelink et al., 2015).

In human cells, B56 isoforms are encoded by five separate genes (B56α, β, γ, δ and ε). The interaction interfaces involved in BubR1 and Sgo1 binding are extremely well conserved between all of these B56 isoforms (supp.fig.1). This explains why BubR1 and Sgo1 appear to display no specificity for individual B56 isoforms (Kitajima et al., 2006; Kruse et al., 2013; Riedel et al., 2006; Xu et al., 2013; Xu et al., 2009), and why these isoforms have been proposed to function redundantly at kinetochores during mitosis (Foley et al., 2011).

However, one crucial observation throws doubt over this issue of redundancy: individual B56 isoforms localise differentially to either the kinetochore or centromere in human cells (Meppelink et al., 2015; Nijenhuis et al., 2014). It is therefore not easy to reconcile this differential localisation with the evidence presented above, which implies that the centromere and kinetochore receptors for B56 do not display any selectivity for individual isoforms. This caused us to readdress the question of redundancy and isoform specificity in human cells.

## RESULTS

### PP2A-B56 isoforms have specific roles at the kinetochore during mitosis

PP2A-B56 isoform localisation to the centromere and kinetochore was visualised in nocodazole-arrested HeLa Flp-in cells expressing YFP-tagged B56 subunits. This revealed that while some B56 isoforms localise to the centromere (B56α and ε), others localise to the outer kinetochore (B56γ and δ), and one isoform displayed a mixed localisation pattern (B56β) (Figures 1a,b). B56 isoforms have been proposed to act redundantly at the kinetochore in human cells (Foley et al., 2011), therefore we readdressed this question in light of their differential localisation. We chose to compare B56α and B56γ as representative members of the centromere and kinetochore-localised pools, respectively. We first confirmed that endogenously tagged YFP-B56α and YFP-B56γ displayed the same differential localisation to the centromere or kinetochore (supp.fig.2). We then knocked down all B56 isoforms, except for either B56α or B56γ (supp.fig.3), to determine whether these endogenous isoforms could support kinetochore functions. As expected (Espert et al., 2014; Nijenhuis et al., 2014), simultaneous depletion of all B56 isoforms enhanced basal Knl1-MELT phosphorylation, delayed MELT dephosphorylation upon Mps1 inhibition with AZ-3146 (Hewitt et al., 2010), and prevented mitotic exit under identical conditions (Figure 1c-e). Importantly, these effects were all rescued when endogenous B56γ was preserved, but not if only B56α remained (Figure 1c-e). Kinetochore PP2A-B56 also has well-established roles in chromosome alignment where it is needed to antagonise Aurora B and allow initial kinetochore-microtubule attachment to form (Foley et al., 2011; Kruse et al., 2013; Suijkerbuijk et al., 2012; Xu et al., 2013). Knockdown of all B56 isoforms produced severe chromosome alignment defects, which could be rescued by preserving B56γ, but not B56α (Figure 1f). In summary, only the kinetochore-localised B56γ, and not the centromeric B56α, can support SAC silencing and chromosome alignment in human cells.

**Fig 1:**
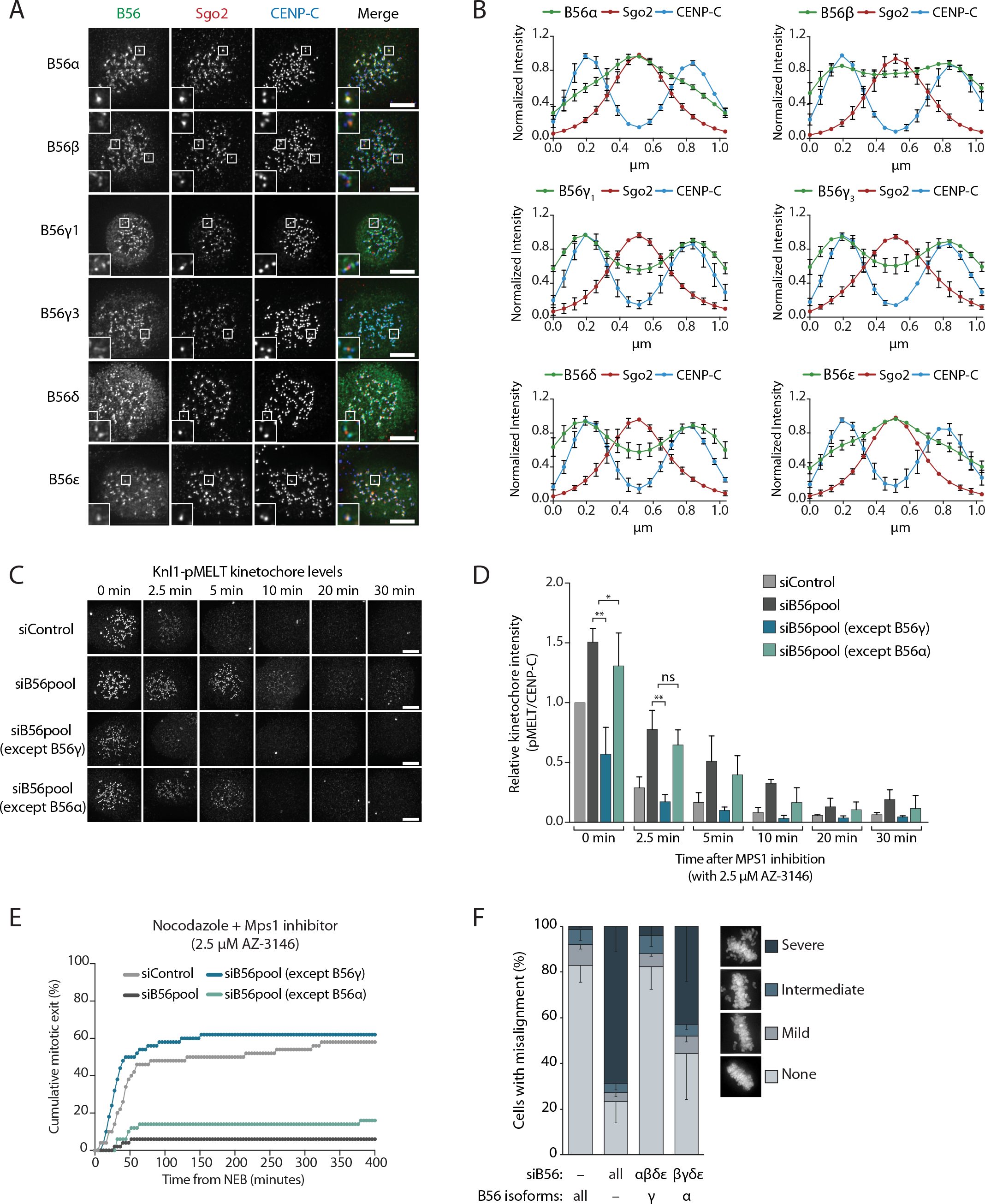
A subset of PP2A-B56 complexes control spindle assembly checkpoint silencing and chromosome alignment. **A** and **B.** Representative images (**A**) and line plots (**B**) of nocodazole-arrested Flp-in HeLa cells expressing YFP-B56 (B56α, B56β, B56γ1, B56γ3, B56δ and B56ε). For line plots, 5 kinetochore pairs were analysed per cell, for a total of 10 cells per experiment. Graphs represent the mean intensities (+/− SD) from 3 independent experiments. Intensity is normalized to the maximum signal in each channel in each experiment. **C-F.** Flp-in HeLa cells treated with siRNA against B56pool, all B56 isoforms except B56α, or all B56 isoforms except B56γ were analysed for SAC silencing and chromosomal alignment. Representative images (**C**) and quantification (**D**) of relative kinetochore intensities of Knl1-pMELT in cells arrested in prometaphase with nocodazole and treated with MG132 for 30 minutes, followed by 2.5 µM AZ-3146 for the indicated amount of time. 10 cells were quantified per experiment and the graph represents the mean (+ SD) of 3 independent experiments. **E.** Time-lapse analysis of cells entering mitosis in the presence of nocodazole and 2.5 µM AZ-3146. The graph represents the cumulative data from 50 cells, which is representative of 3 independent experiments. **F.** Quantification of chromosome misalignment in cells arrested in metaphase with MG-132. At least 100 cells were scored per condition per experiment and graph represents the mean (-SD) of 3 independent experiments. Misalignments were score as mild (1 to 2 misaligned chromosomes), intermediate (3 to 5 misaligned chromosomes), and severe (>6 misaligned chromosomes). Asterisks indicate significance (Welch’s t-test, unpaired); ns P > 0.05, * P ≤ 0.05, ** P < 0.01.

Overexpression of GFP-B56α has previously been shown to rescue kinetochore-microtubule attachment defects following the depletion of all PP2A-B56 isoforms in human cells (Foley et al., 2011). To understand the discrepancy with our data, we performed the same assays as previously, but this time expressing siRNA-resistant YFP-B56α or YFP-B56γ to rescue the knockdown of all endogenous B56 isoforms. Under these conditions, both exogenous B56 isoforms were able to rescue MELT dephosphorylation, SAC silencing and chromosome alignment (supp.fig.4). The ability of exogenous YFP-B56α to support kinetochore functions can be explained by the fact that it is highly overexpressed, which leads to elevated centromere and kinetochore levels in comparison to the endogenous YFP-B56α situation (supp.fig.5). We therefore conclude B56α acts primarily at the centromere, but it can still function at the kinetochore when overexpressed.

### Sgo2 provides specificity for centromeric B56 recruitment

We next sought to determine the reason for differential B56 isoform localisation, which was puzzling because the reported kinetochore and centromere receptors - BubR1 and Sgo1 - do not appear to display selectivity for individual B56 isoforms (Kitajima et al., 2006; Kruse et al., 2013; Riedel et al., 2006; Xu et al., 2013; Xu et al., 2009). We initially focussed on the centromere receptor because Sgo1 and Sgo2 can both bind to PP2A-B56 (Rivera et al., 2012; Tanno et al., 2010; Xu et al., 2009). Sgo1 is considered the primary receptor because it is more important than Sgo2 for protecting cohesion in mitosis (Huang et al., 2007; Kitajima et al., 2005; Kitajima et al., 2006; Llano et al., 2008; McGuinness et al., 2005; Rivera et al., 2012; Tang et al., 2006; Tanno et al., 2010), but this could be explained by both PP2A dependent and independent effects that are specific to Sgo1 (Hara et al., 2014; Kitajima et al., 2006; Liu et al., 2013b; Nishiyama et al., 2013). In fact, the only study that has directly compared the contribution of Sgo1 and Sgo2 to centromeric PP2A-B56 recruitment, has concluded that Sgo2 is more important (Kitajima et al., 2006). We therefore set out to clarify the role of Sgo1 and Sgo2 in controlling the recruitment of B56 isoforms to the centromere in human cells.

Depletion of Sgo2, but not Sgo1, caused a significant reduction in B56α levels at the centromere (Figures 2a-d). Although Sgo1 depletion did not reduce total centromeric B56α, it did cause both Sgo2 and B56α to spread out from the centromere towards the kinetochore (figure 2e), as shown previously by others (Meppelink et al., 2015). This is due to inefficient anchoring of Sgo2 at centromeres because combined Sgo1 and Sgo2 depletion completely removed B56α from kinetochores/centromeres (figure 2f,g). We therefore conclude that, as suggested previously by others (Kitajima et al., 2006), Sgo2 is the primary centromeric receptor for PP2A-B56 during mitosis. However, Sgo1 also contributes to centromeric B56 localisation primarily by helping to anchor the Sgo2-B56 complex at the centromere, perhaps by bridging an interaction with cohesin rings (Hara et al., 2014; Liu et al., 2013b).

**Figure 2:**
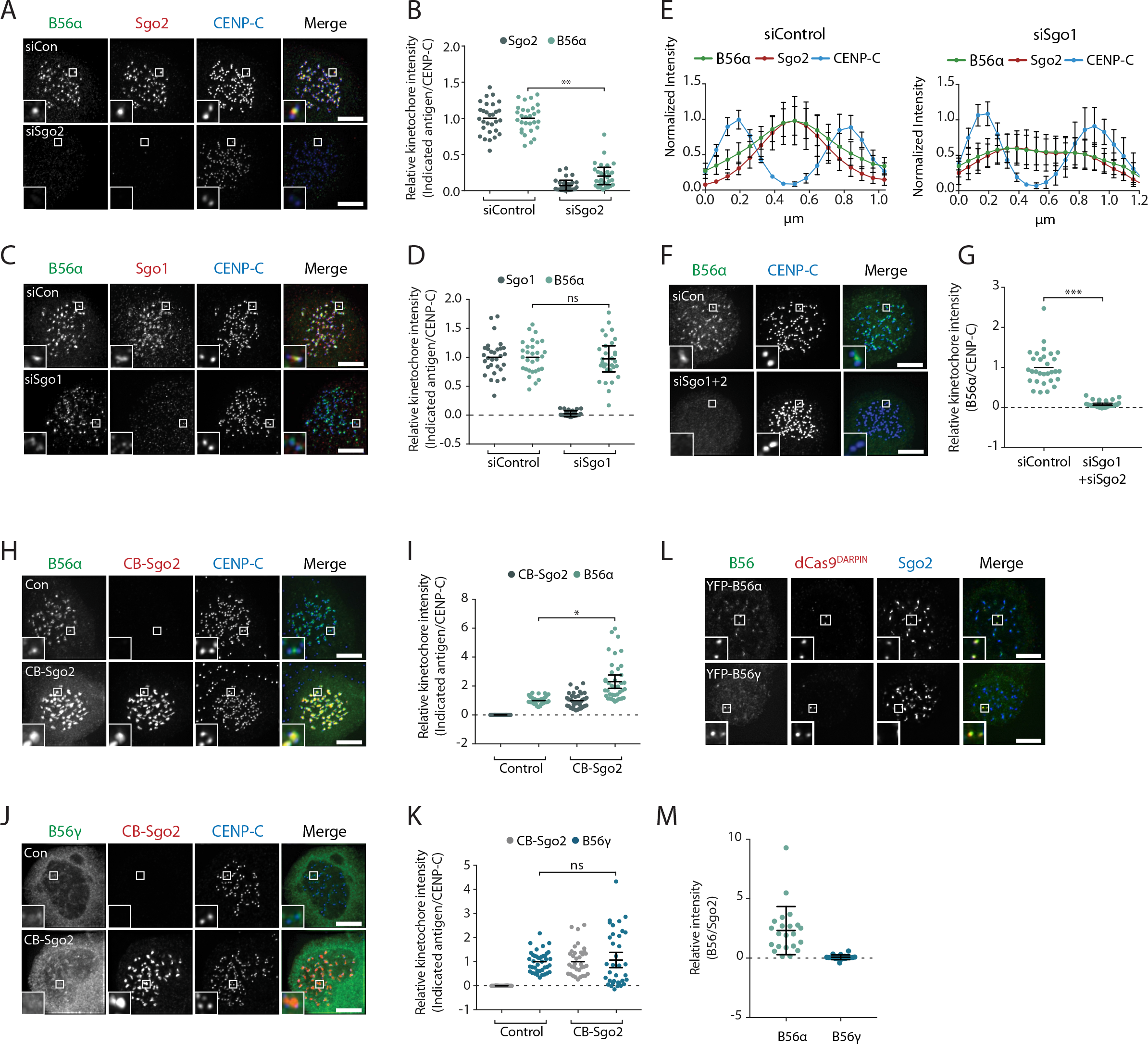
Sgo2 specifically localizes B56α to centromeres. **A-G.** The effect of Sgo1 and/or Sgo2 knockdown on YFP-B56α localisation in Flp-in HeLa cells. Representative images (**A**, **C**, **F**) and quantifications (**B, D**, **G**) of relative kinetochore intensity of B56α in cells arrested in prometaphase with nocodazole after knockdown of Sgo2 (**A**, **B**), Sgo1 (**C**, **D**), or Sgo1 + Sgo2 (**F, G**). **E** shows line plots of Sgo2 and B56α localisation following Sgo1 knockdown; 5 kinetochore pairs were analysed per cell, for a total of 10 cells per experiment. Graphs represent the mean intensities (+/− SD) from 3 independent experiments. Intensity is normalized to the maximum signal present in each channel within the endogenous B56α experiment. **H-M.** Flp-in HeLa cells expressing YFP-B56α or YFP-B56γ were transfected with the CB-Sgo2 (**H-K**) or gChr7+Cas9-DARPIN (**L**, **M**) and analysed for B56 recruitment in cells arrested in prometaphase with nocodazole. **H**, **L**, and **J**. are representative images; **I**and **K.** are quantifications of relative kinetochore intensity of the indicated antigen; and **M** is quantification of intensity of Sgo2 over B56 at the Chr7 locus. For all kinetochore intensity graphs, 10 cells were quantified per experiment and graphs represent the mean (+/− SD) of 3 independent experiments. Asterisks indicate significance (Welch’s t-test, unpaired); ns P > 0.05, * P ≤ 0.05, ** P < 0.01.

We next examined whether specific binding to Sgo1 and/or Sgo2 could explain differential B56 isoform localisation. To address this, we artificially relocalized Sgo1 or Sgo2 to the inner kinetochore, by fusing it to the kinetochore-targeting domain of CENP-B (CB). This location was chosen because it could be distinguished from the endogenous centromeric B56 pool, and yet should still be accessible to Aurora B. This may be important because phosphorylation of Sgo2 by Aurora B has been proposed to be needed for B56 interaction (Tanno et al., 2010). Whereas CB-Sgo1 was able to localise both B56α and B56γ to the inner kinetochore (supp.fig.6), CB-Sgo2 was only able to recruit B56α (figure 2h-k). To confirm that endogenous Sgo2 displayed selectivity for specific B56 isoforms, we used a Designed Ankyrin Repeat Protein (DARPin) that can bind to GFP with high affinity (Brauchle et al., 2014). The DARPin was fused to dCas9 to enable the selective targeting of YFP-tagged B56α or B56γ to a repetitive region on chromosome 7 (Chr7). This assay confirmed that only B56α, and not B56γ, was able to co-recruit endogenous Sgo2 to this region (figure 2l, m). Considering Sgo2 is the primary centromeric receptor for B56 (figure 2a, b) (Kitajima et al., 2006), this provides an explanation for why only a subset of B56 isoforms localise to the centromere.

### Sgo1 collaborates with BubR1 to recruit B56 to kinetochores

We next turned our attention to the reason for differential kinetochore localisation. PP2A-B56 binds to kinetochores by interacting with a phosphorylated LxxIxE motif in BubR1 (Kruse et al., 2013; Suijkerbuijk et al., 2012; Xu et al., 2013) and this interaction is mediated by a binding pocket on B56 that is completely conserved in all isoforms (supp.fig.1) (Hertz et al., 2016; Wang et al., 2016a; Wang et al., 2016b). Therefore, we hypothesised that additional interactions may help to stabilise specific B56 isoforms at the kinetochore. In agreement with this hypothesis, BubR1 depletion and/or mutation of the LxxIxE binding pocket in B56γ (B56γ ^H187A^) reduced but did not completely remove B56γ from kinetochores (figure 3a-d, supp. fig 7a, b). The remaining B56γ in these situations spreads out between the kinetochore and centromere (figure 3e,f), which implies that B56γ uses additional interactions to be maintained at centromeres/kinetochores.

**Figure 3:**
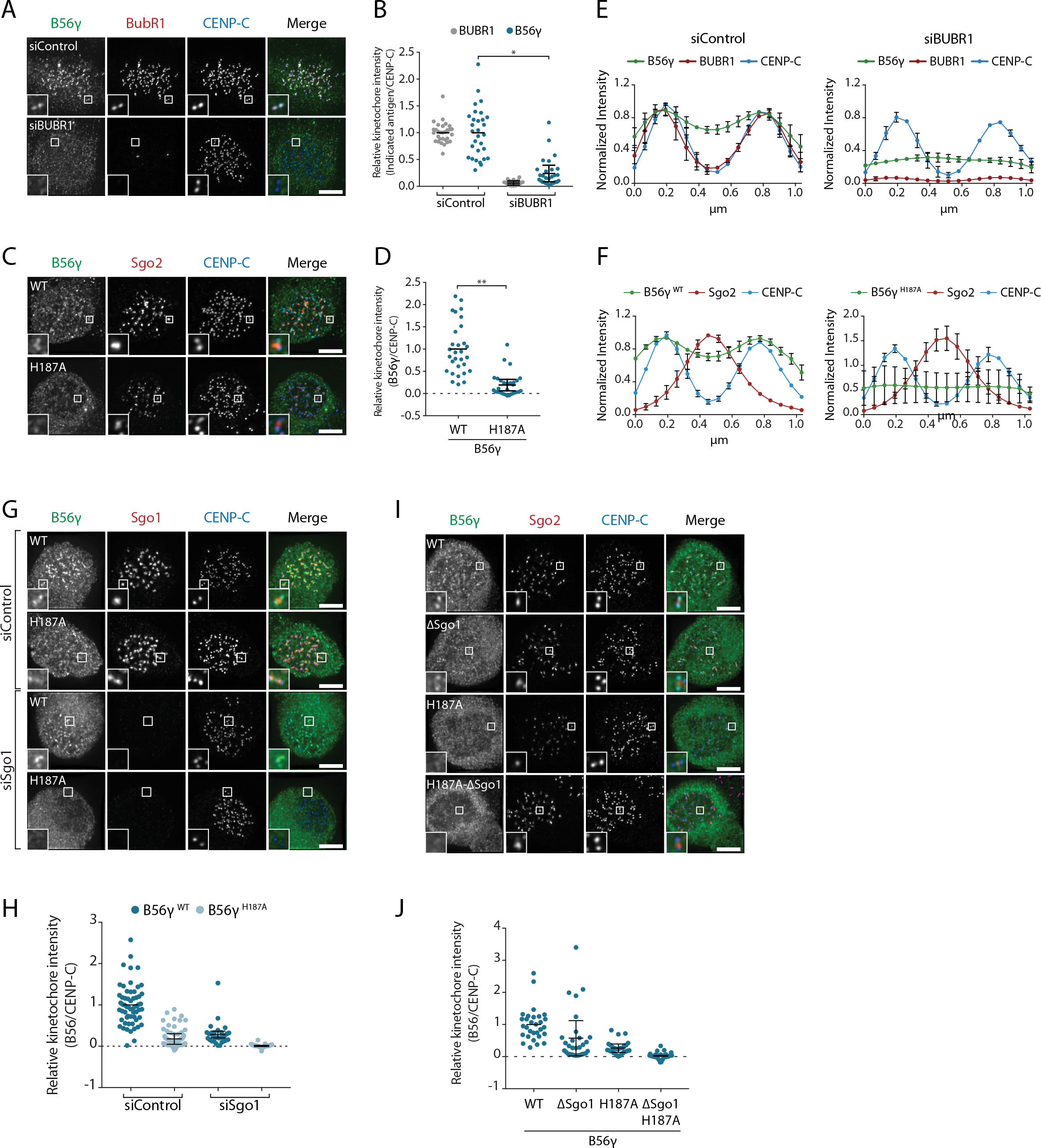
BubR1 and Sgo1 localize B56γ to kinetochores. B56γ kinetochore localisation in Flp-in HeLa cells after BubR1 knockdown (**A**, **B**, **E**) or mutation of the LxxIxE binding pocket (H187A: **C**, **D**, **F**) in cells arrested in prometaphase with nocodazole. For each condition, representative images (**A**, **C**), quantification of relative kinetochore levels (**B**, **D**) and line plot analysis (**E**, **F**) depicts the levels and distribution of the indicated antigens. **G-J**: representative images (**G**, **I**) and quantification of relative kinetochore intensities (**H**, **J**) YFP-B56γ WT or H187A following Sgo1 knockdown (G, H) of mutation of the Sgo1 binding region (ΔSgo1). Each graph represents the mean intensities (+/−SD) from at least 3 independent experiments. For the line plot analysis, 5 kinetochore pairs were analysed per cell, for a total of 10 cells per experiment. Intensity is normalized to the maximum signal in each channel in each experiment. Asterisks indicate significance (Welch’s t-test, unpaired); ns P > 0.05, * P ≤ 0.05, ** P < 0.01.

A targeted siRNA screen identified critical roles for Knl1 and Bub1, which, when depleted, completely abolished B56γ recruitment to kinetochores (supp.fig.7c-f). Knl1 recruits Bub1 to kinetochores, and Bub1 scaffolds the recruitment of BubR1 (Johnson et al., 2004; Overlack et al., 2015; Primorac et al., 2013). However, in addition to this, Bub1 also phosphorylates histone H2A to localise Sgo1 to histone tails that are adjacent to the kinetochore (Baron et al., 2016; Kawashima et al., 2010; Kitajima et al., 2005; Liu et al., 2013a; Tang et al., 2004; Yamagishi et al., 2010). Since Sgo1 can bind to B56γ (supp.fig.6) we examined its role in the kinetochore recruitment of this isoform. Sgo1 depletion reduced B56γ ^WT^ at kinetochores and completely removed B56γ ^H187A^ (figure 3g,h). Moreover, this was specific for Sgo1, because Sgo2 depletion had no effect (supp.fig.7g, h). To test whether this was due to direct binding to Sgo1 we generated a B56 Sgo1-binding mutant (B56γ^ΔSgo1^), which we confirmed was defective in binding CB-Sgo1 *in vivo* (supp.fig.8). This mutation reduced the recruitment of B56γ WT to kinetochores and completely abolished the recruitment of B56γ ^H187A^ (figure 3i,j), in a manner that was similar to the effect of Sgo1 depletion (figure 3g,h). This demonstrates that Bub1 establishes two separate arms that cooperate to recruit B56γ to kinetochores: it binds directly to BubR1, which interacts via its LxxIxE motif with B56γ, and it phosphorylates Histone-H2A to recruit Sgo1, which additionally helps to anchor B56γ at kinetochores.

### B56 isoforms bind differentially to LxxIxE containing motifs during mitosis

The B56-Sgo1 interaction is unlikely to explain B56 isoform specificity at kinetochores, since Sgo1 interacts with both B56α and B56γ when recruited to centromeres (supp.fig.6). We therefore focussed on the LxxIxE interaction with BubR1 to quantitatively assess the binding to B56α and B56γ. Immunoprecipitations of equal amounts of B56α and B56γ from nocodazole-arrested cells demonstrated that BubR1 bound preferentially to B56γ (figure 4a,b). Moreover, a panel of antibodies against other LxxIxE containing proteins (Hertz et al., 2016), demonstrated that LxxIxE binding was generally reduced in B56α immunoprecipitates (figure 4a,b). B56γ has been shown to display slightly higher affinities for some LxxIxE containing peptides *in vitro* (Wu et al., 2017), which in principle could allow this isoform to outcompete B56α for binding. However, a simple competition model is unlikely to explain differential kinetochore localisation, since we observe no change in B56α localisation when all other B56 isoforms are present or knocked down (Figure 4c,d). Instead, we favour the hypothesis that binding to LxxIxE motifs is specifically perturbed in PP2A-B56α complexes during prometaphase.

**Figure 4:**
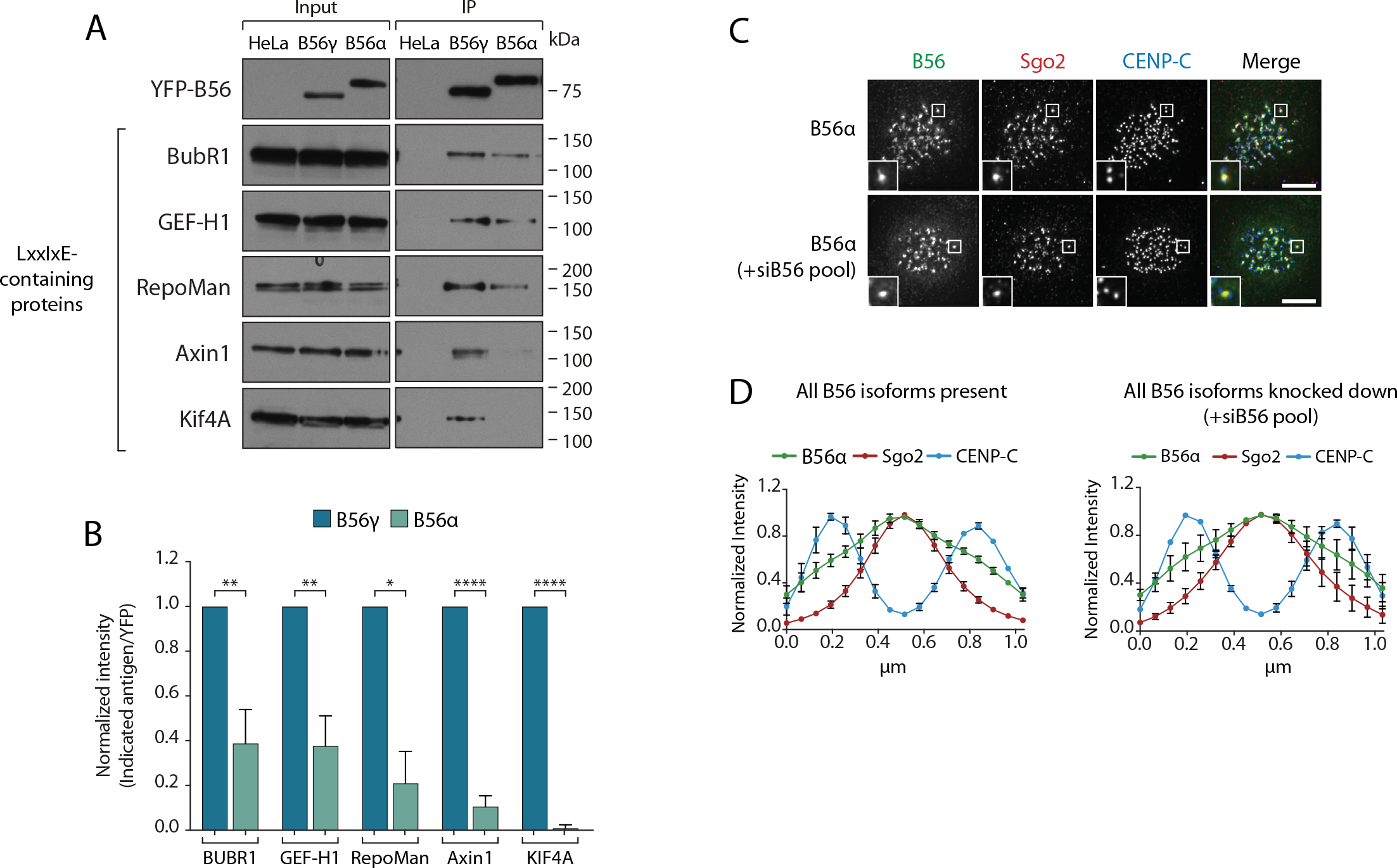
Specific binding of B56γ to kinetochores reflects an enhanced ability to bind LxxIxE motifs. **A.**Immunoblot of the indicated proteins, containing a LxxIxE motif (Hertz et al., 2016), following YFP immunoprecipitation from nocodazole-arrested Flp-in HeLa cells expressing YFP-B56α or YFP-B56γ. **B.** Quantification of the mean normalised intensity (+SD) of the indicated antigens in B56α immunoprecipitates, relative to B56γ immunoprecipitates, from at least 3 experiments. Representative images (**C**) and line plot analysis (**D**) of YFP-B56α in Flp-in HeLa cells arrested in nocodazole and treated with the indicated siRNA. Each line plot graph represents the mean intensities (+/− SD) from 3 independent experiments. 5 kinetochore pairs were analysed per cell, for a total of 10 cells per experiment. Intensity is normalized to the maximum signal in each channel in each experiment. Asterisks indicate significance (Welch’s t-test, unpaired); ns P > 0.05, * P ≤ 0.05, ** P < 0.01, *** P < 0.001, **** P < 0.0001.

### Residues within a C-terminal loop of B56 determine localisation to the centromere or kinetochore

We next searched for the molecular explanation for differential B56 isoform localisation. To do this, we generated four chimaeras between B56α and B56γ by joining the isoforms in the loops that connect the α-helixes (figure 5a). Immunofluorescence analysis demonstrated that B56γ localisation switched from kinetochores to centromeres in chimaera 4 (figure 5b,c). Furthermore, this region alone is sufficient to switch localisation to the centromere when transferred into B56γ, and the corresponding region in B56γ can induce localisation to the kinetochore if transplanted into B56α (supp fig.9). We generated four additional chimaeras to narrow down this region even further to amino acids 405-425 in B56α, which contains an α-helix and a small loop that juxtaposes the catalytic domain in the PP2A-B56 complex (figure 5d-f). Importantly, switching just 4 amino acids within this loop in B56α to the corresponding residues in B56γ (B56α ^TKHG^) was sufficient to relocalise B56α from centromeres to kinetochores (figure 5g-i). Furthermore, the B56α ^TKHG^ remained functional and holoenzyme assembly was unperturbed (supp fig.10). In summary, a small C-terminal loop in B56 defines whether B56 localises to centromeres, via Sgo2, or to kinetochores, via an LxxIxE interaction with BubR1.

**Figure 5:**
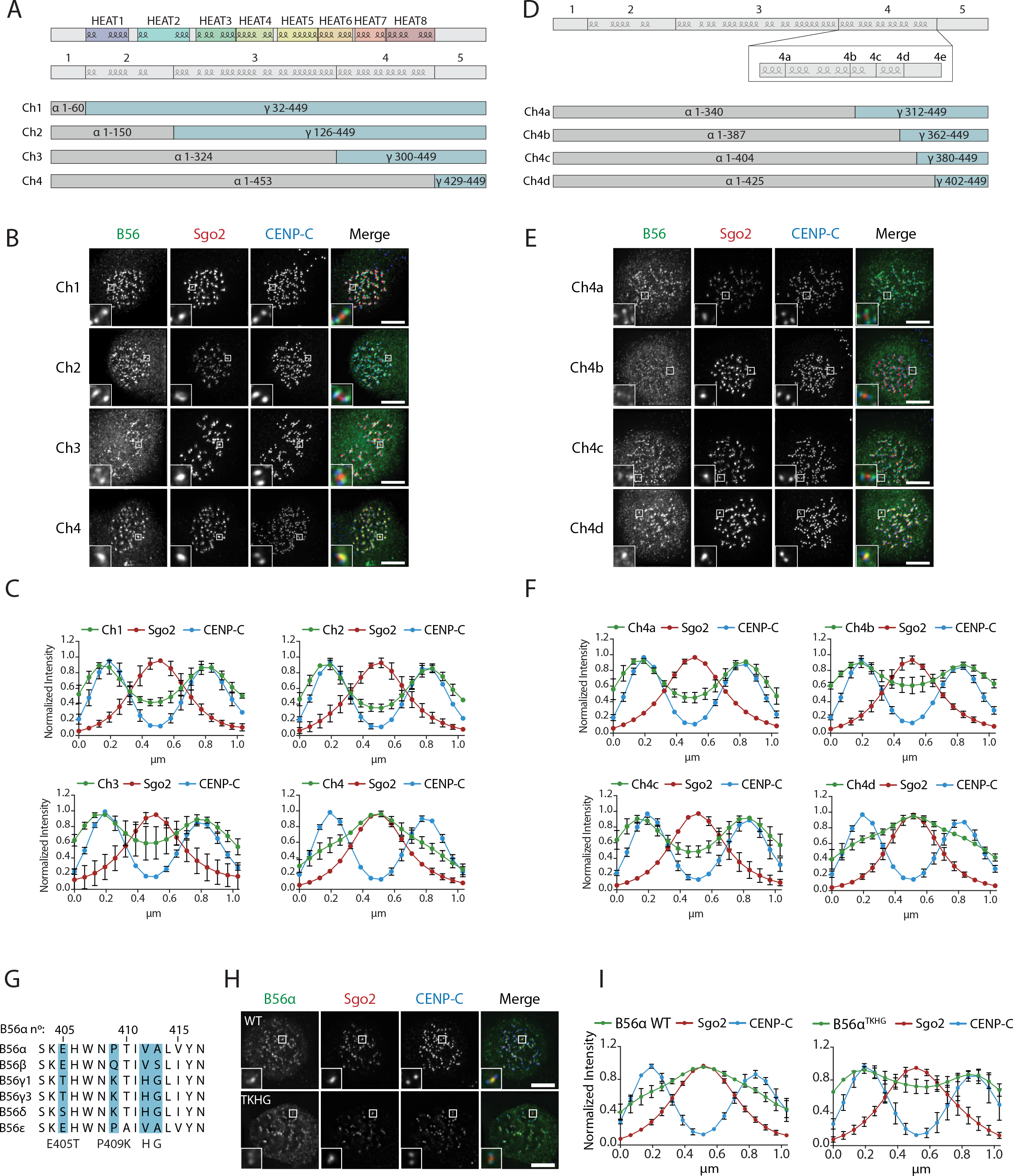
A C-terminal loop in B56 specifies B56 localization to centromeres or kinetochores. B56 localisation in B56α-γ chimeras spanning the entire B56 (Ch1-4: **A-C**), a region at the C-terminus (Ch4a-4d: **D-F**). **A**, **D.**Schematic representation of the B56α-γ chimeras created. Representative images (**B**, **E**) and line plot analysis (**C**, **F**) to show the B56 localisation pattern in each chimaera. **G**. Alignment of B56 isoforms within region 4d that controls centromere/kinetochore localisation. **G-H**: Effect of four point-mutations within region 4d to convert B56α to the correspond B56γ sequence (B56α^TKHG^). Representative images (**H**) and line plot analysis (**I**) of B56α WT or B56α^TKHG^ in cells arrested in prometaphase with nocodazole. Each graph represents the mean intensities (+/− SD) from 3 independent experiments. 5 kinetochore pairs were analysed per cell, for a total of 10 cells per experiment. Intensity is normalized to the maximum signal in each channel in each experiment.

### The C-terminal loop controls Sgo2 binding and LxxIxE motif affinity

We next addressed whether the B56α ^TKHG^ mutant switched the Sgo2 and LxxIxE binding properties of B56α. In-cell interaction assays confirmed that B56α ^TKHG^ was unable to bind to endogenous or exogenous Sgo2 (figure 6a-c). Furthermore, immunoprecipitation of YFP-B56α ^TKHG^ showed an enhanced ability to bind LxxIxE containing proteins and, in particular, BubR1 (figure 6e,f). Therefore, we conclude that the small EPVA loop in B56α is necessary for the interaction with Sgo2 and the centromere and, in addition, it is also required to fully repress binding to LxxIxE motifs and the kinetochore. Importantly, this loop is not sufficient to induce either of these effects when transplanted alone into B56γ, because B56γ ^EPVA^ is not lost from the kinetochore or gained at the centromere (supp.fig.11a). Instead, a region immediately C-terminal to the EPVA (amino acids 414-453 in B56α) is also required to induce centromere binding, and a small helix N-terminal to the EPVA (amino acids 374-386 in B56α) is needed to repress kinetochore binding (supp.fig.11b). Therefore, although the regions that define centromere and kinetochore localisation overlap at the EPVA loop, they have different distal requirements that demonstrate that they are not identical (supp.fig.11c).

**Figure 6:**
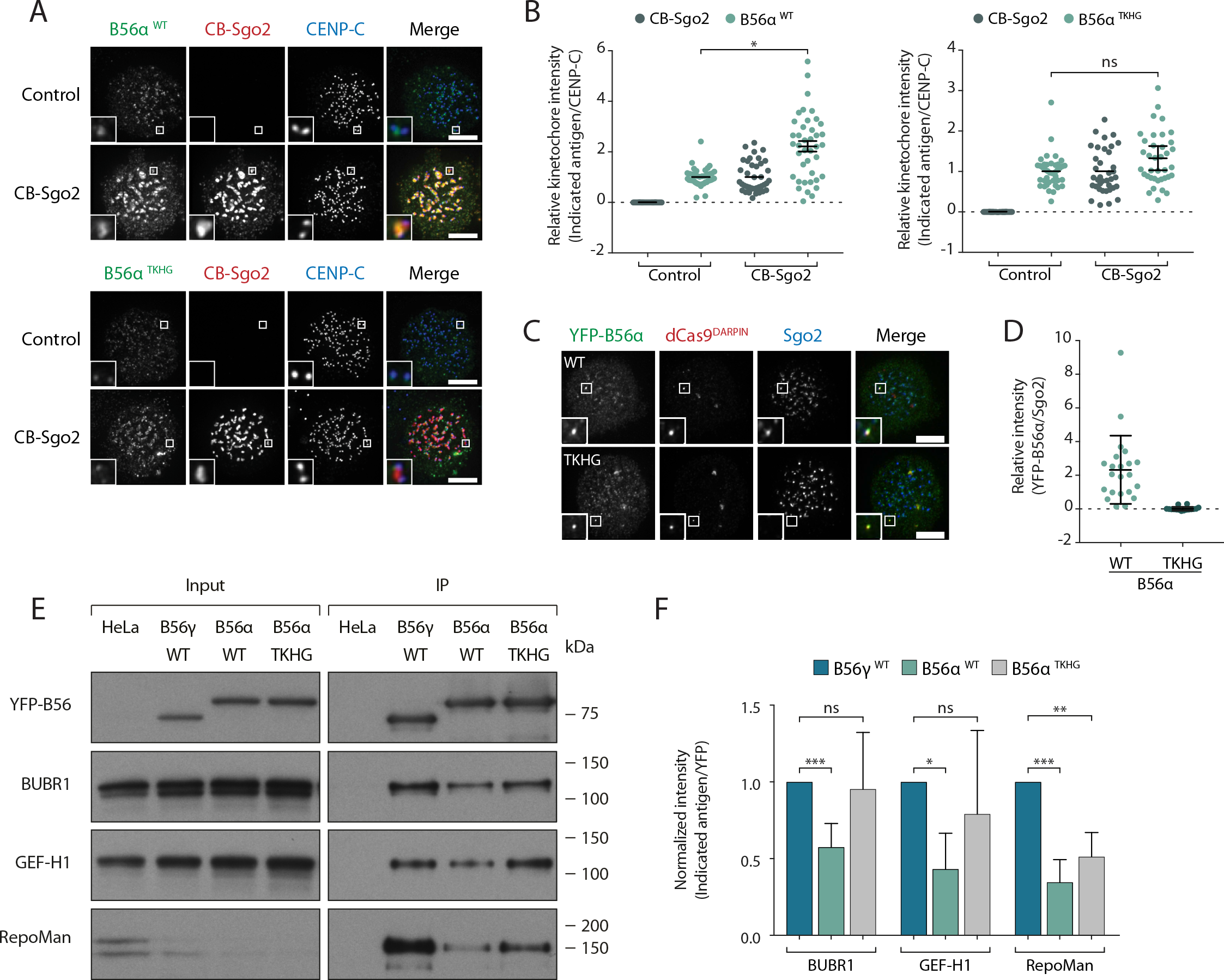
A C-terminal loop in B56 regulates binding to Sgo2 and LxxIxE motifs. **A-D**: Flp-in HeLa cells expressing either YFP-B56α WT or TKHG were transfected with the CB-Sgo2 (**A, B**) or gChr7 + dCas9-DARPIN (**C, D**) and analysed for B56 recruitment. Representative images (**A**, **C**) and quantification of relative kinetochore intensity (**B**) or intensity of Sgo2 over B56α at the Chr7 locus (**D**). For all kinetochore intensity graphs, 10 cells were quantified per experiment and graphs represent the mean (+/− SD) of at least 3 independent experiments. **E.**Immunoblot of the indicated antigens following immunoprecipitation of YFP from nocodazole-arrested Flp-in HeLa cells expressing YFP-B56γ, YFP-B56α WT or YFP-B56α-TKHG. **F.**Quantification of the mean normalised intensity (+SD) of indicated antigens in B56α WT or B56α TKHG immunoprecipitates, relative to B56γ, from at least 4 experiments. Asterisks indicate significance (Welch’s t-test, unpaired); ns P > 0.05, * P ≤ 0.05, ** P < 0.01, *** P < 0.001.

## DISCUSSION

This work demonstrates how different B56 isoforms localise to discrete subcellular compartments and control separate processes during mitosis. Differential B56 isoform localisation has previously been observed in interphase (McCright et al., 1996) and during the later stages of mitosis (Bastos et al., 2014), which implies that B56 isoforms may have evolved to carry out specific functions, at least in part, by targeting PP2A to distinct subcellular compartments. The differential localisation we observe during prometaphase arises because B56 isoforms display selectivity for specific receptors at the centromere and kinetochore.

The centromeric isoform B56α binds preferentially to Sgo2 via a C-terminal stretch that lies between amino acids 405 and 453. A key loop within this region juxtaposes the catalytic domain and contains an important EPVA signature that is critical for Sgo2 binding and is unique to B56α and B56ε. This sequence is also conserved in *Xenopus* B56ε, which has previously been shown to selectively bind to Sgo2, when compared to B56γ (Rivera et al., 2012). We therefore propose that a subset of B56 isoforms (B56α and ε) utilize unique motifs to interact with Sgo2 and the centromere during mitosis. How then, can these results be reconciled with the fact that Sgo1 appears to be more important than Sgo2 for the maintenance of cohesion during mitosis (Huang et al., 2007; Kitajima et al., 2005; Kitajima et al., 2006; Llano et al., 2008; McGuinness et al., 2005; Rivera et al., 2012; Tang et al., 2004; Tang et al., 2006; Tanno et al., 2010)? One possibility is that this reflects a dual role for Sgo1 in both preserving Sgo2-PP2A-B56 at centromeres and competing with the cohesin release factor, WAPL, for cohesin binding (Hara et al., 2014). Alternatively, perhaps only Sgo1-PP2A-B56 complexes are able to preserve cohesion because Sgo1 is able to bind to SA2–Scc1 directly (Hara et al., 2014; Liu et al., 2013b; Tanno et al., 2010), thereby positioning PP2A-B56 to dephosphorylate nearby residues within the cohesin complex. In that case, the small amount of Sgo1-PP2A-B56α/ε that remains at centromeres following Sgo2 depletion (figure 2a,b) could be sufficient to preserve cohesion.

The kinetochore B56 isoforms bind to BubR1 via a canonical LxxIxE motif within the KARD (Hertz et al., 2016; Kruse et al., 2013; Suijkerbuijk et al., 2012; Xu et al., 2013). Although the LxxIxE binding pocket is completely conserved in all B56 isoforms (supp.fig.1), we observe a striking preference in the binding of B56γ over B56α to many LxxIxE containing proteins during prometaphase (fig. 4). We hypothesise that this is due to repressed binding between LxxIxE motifs and B56α during prometaphase, because LxxIxE binding (fig.6e,f) and kinetochore accumulation (figure 5h,j) can both be enhanced by mutation of the EPVA loop in B56α (B56α ^TKHG^). We cannot, however, exclude the possibility that the corresponding TKHG sequence in B56γ positively regulates LxxIxE interaction and kinetochore localisation. Considering that this region also controls Sgo2 and centromere binding, a simple explanation could be that Sgo2 interaction obscures the LxxIxE binding pocket; however, this appears unlikely given that Sgo2 depletion does not relocalise B56α to kinetochores (figures 2a,b). Furthermore, centromere and kinetochore binding can occur together and the regions that define each of these localisations do not fully overlap (supp.fig.11). Instead, we speculate that another interacting partner, or alternatively a tail region within a PP2A-B56 subunit, might obscure or modify the conformation of the LxxIxE binding pocket in B56α.

An important additional finding of this work is that Sgo1 contributes to the retention of B56 isoforms at both the centromere and the kinetochore: Sgo1 depletion reduces B56γ levels at the kinetochore (figure 3g,h) and causes B56α to spread out from the centromere (figure 2e). Furthermore, if Sgo2 or BubR1 is depleted to inhibit B56 localisation to centromeres or kinetochores, then, in both cases, the B56 isoforms that remain are bound to Sgo1 and spread out along the centromere-kinetochore axis (figure 3 and results not shown). It will be important in future to determine exactly how Sgo1 collaborates with BubR1 and Sgo2 to control B56 localisation and, in particular, whether Sgo1 can interact with Sgo2-B56 or BubR1-B56 complexes directly, or whether this is prevented by mutually exclusive interactions. The interfaces between BubR1-B56 and Sgo1-B56 do not appear to be overlapping, at least based on current structural data (Wang et al., 2016a; Wang et al., 2016b; Xu et al., 2009), which implies that Knl1-bound BubR1-B56 could potentially be anchored towards histone tails by Sgo1. This could have important implications for SAC signalling and tension-sensing.

In summary, the work presented here demonstrates how different members of the PP2A-B56 family function during the same stage of mitosis to control different biological processes. This is the first time that such sub-functionalisation has been demonstrated between isoforms of the same B family. It is currently unclear why such specialisation is necessary or at least preferable to a situation whereby all B56 isoforms operate redundantly, as initially suggested (Foley et al., 2011). One possibility is that the use of different B56 isoforms allows PP2A catalytic activity to be regulated differently in specific subcellular compartments: for example, by interactions or post-translational modifications that are specific for the B56 subunits. In this respect, protein inhibitors of PP2A-B56 have been shown to function specifically at the centromere (SET (Chambon et al., 2013)) and at the kinetochore (BOD1 (Porter et al., 2013)); therefore, it would be interesting to test whether these inhibitors display selectivity for certain PP2A-B56 isoforms. Future studies such as this, which build upon the work presented here, may ultimately help to reveal novel ways to modulate the activity of specific PP2A-B56 complexes. The recent development of selective inhibitors of related PP1 regulatory isoforms to combat neurodegenerative diseases (Das et al., 2015; Krzyzosiak et al., 2018), provides a proof-of-concept that successful targeting of specific phosphatase isoforms is both achievable and therapeutically valuable.

## ACKNOWLEDGEMENTS

This work was funded by Cancer Research UK (C47320/A21229 to ATS) and Tenovus Scotland. We thank staff at the Dundee Imaging Facility and the Flow Cytometry and Cell Sorting facility. We also thank Prof Stephen Taylor for providing the HeLa Flp-in cell line, Timothy Yen and Susanne Lens for providing plasmids, and Iain Cheeseman for useful discussions.

## METHODS

### Cell culture and reagents

HeLa Flp-in cells (Tighe et al., 2008), stably expressing a TetR, were cultured in DMEM supplemented with 9% tetracycline-free FBS, 50 μg/mL penicillin/streptomycin and 2 mM L-glutamine. All cell lines were routinely screened (every 4–8 weeks) to ensure they were free from mycoplasma contamination. All HeLa Flp-in cells stably expressing a doxycycline-inducible construct were derived from the HeLa Flp-in cell line by transfection with the pCDNA5/FRT/TO vector (Invitrogen) and the FLP recombinase, pOG44 (Invitrogen), and cultured in the same medium but containing 200 μg/mL hygromycin-B. Plasmids were transfected using Fugene HD (Promega) according to manufacturer’s protocol.

1 µg/mL doxycycline was added for ≥16 h to induce protein expression in the inducible cell lines. Thymidine (2 mM) and nocodazole (3.3 µM) were purchased from Millipore, MG132 (10 µM) and AZ-3146 from Selleck Chemicals, doxycycline (1 µg/mL) from Sigma, 4,6-diamidino-2-phenylindole (DAPI, 1:50000) from Invitrogen, AZ-3146 from Axon, calyculin A (10 µM in 10% EtOH) from LC labs, RO-3306 (10 µM) from Tocris and hygromycin-B from Santa Cruz Biotechnology.

### Plasmids and cloning

pCDNA5-YFP-B56α, β, γ1, γ3, δ and ε were amplified from pCEP-4xHA-B56 (Addgene plasmids 14532-14537; deposited by D. Virshup, Duke-NUS Graduate Medical School, Singapore) and subcloned into pCDNA5-LAP-BubR1^WT^ (Nijenhuis et al., 2014) through Not1 and Apa1 restriction sites. B56γ1 and B56γ3 were corrected to start on M1 and not 11, and the R494L mutation in B56γ3 was corrected. pCDNA5-YFP-B56α and pCDNA5-YFP-PP2A-B56γ1 were made siRNA-resistant by site-directed mutagenesis (silent mutations in the coding sequence for E102 and L103 in B56α, and T126 and L127 in B56γ). All B56α and B56γ1 mutants were created by site-directed mutagenesis from pCDNA5-YFP-B56α and pCDNA5-YFP-B56γ1, respectively. The B56α–γ chimeras were generated by Gibson assembly with pCDNA5-YFP-B56α and pCDNA5-YFP-B56γ used as templates for the PCR reaction. vsv-CENP-B-Sgo1-mCherry (Meppelink et al., 2015) was used to make vsv-CENP-B-Sgo2-mCherry, by removing Sgo1 and adding Sgo2 via Gibson assembly from pDONR-Sgo2 (a gift from T. J. Yen). The Sgo1 binding mutant in B56γ (B56γ^ΔSgo1^) was created by site directed mutagenesis to create 3 mutations: Y391F, L394S and M398Q. The dCas9-DARPIN-flag was created by digesting pHAGE-TO-dCas9-3xmCherry (Addgene #64108) with BamHI and XhoI to remove 3xmCherry and replace with a synthesised DARPIN-flag that binds to GFP with high affinity (Brauchle et al., 2014). The gRNA targeting a repetitive region on chromosome 7 was generated by PCR mutagenesis to introduce the gRNA sequence (GCTCTTATGGTGAGAGTGT (Chen et al., 2016)) into the pU6 vector.

### Gene knockdowns

Cells were transfected with 20 nM siRNA using Lipofectamine^®^ RNAiMAX Transfection Reagent (Life Technologies) according to the manufacturer’s instructions. For simultaneous knockdown of all B56 isoforms (B56pool) the single B56 isoform siRNA were mixed at equimolar ratio of 20 nM each. The siRNA sequences used in this study are as follows: B56α (PPP2R5A), 5’-UGAAUGAACUGGUUGAGUA-3’; B56β (PPP2R5B), 5’-GAACAAUGAGUAUAUCCUA-3’; B56γ (PPP2R5C), 5’-GGAAGAUGAACCAACGUUA-3’; B56δ (PPP2R5D), 5’-UGACUGAGCCGGUAAUUGU-3’; B56ε (PPP2R5E), 5’-GCACAGCUGGCAUAUUGUA-3’; Sgo1, 5’-GAUGACAGCUCCAGAAAUU-3’; Sgo2, 5’-GCACUACCACUUUGAAUAA-3’; BubR1, 5’-AGAUCCUGGCUAACUGUUC-3’; Knl1, 5’-GCAUGUAUCUCUUAAGGAA-3’; Bub1 5’-GAAUGUAAGCGUUCACGAA-3’; Control (GAPDH), 5’-GUCAACGGAUUUGGUCGUA-3’;. All siRNA oligos were custom made and purchased from Sigma, except for Sgo1, which was ordered from Dharmacon (J-015475-12).

### Expression of B56 isoforms

For reconstitution of B56 isoforms or mutants, HeLa Flp-in cells were transfected with 100nM B56pool or mock siRNA and, in some experiments, 20nM additional control, Sgo1, Sgo2, BubR1, Bub1 or Knl1 siRNA. Cells were transfected with the appropriate siRNA for 16h, after which they were arrested in S phase for 24h by addition of thymidine. Subsequently, cells were released from thymidine for 8–10h and arrested in prometaphase by the addition of nocodazole. YFP-B56 expression was induced by the addition of doxycycline during and following the thymidine block. For BubR1 knockdowns and for all chromosome alignment assays, cells were released from thymidine for 6.5h and arrested at the G2/M boundary with RO3306 for 2h. Cells were then released into nocodazole (BubR1 experiments) or normal growth media (alignment assays) for 15 mins before MG132 was then added for 30 mins to prevent mitotic exit. For alignment assays, this is critical to analyse the synchronous alignment of mitotic cells over a 45-minute period.

### In-cell protein-protein interaction assay using dCas9

Cell were transfected with dCas9-DARPIN-flag and a guide RNA that targets a repetitive region on chromosome 7 (at 1:3 ratio of dCas9:gRNA). Doxycycline was added to induce YFP-B56 isoform expression and 48 h later cells arrested in mitosis were fixed, stained and imaged for co-localisation of YFP-B56 isoforms and Sgo2. Only cells containing defined Flag-dCas9 spots that also co-recruited YFP-B56 were imaged. The majority of these spots recruited YFP-B56, but the dCas9 spots themselves were only readily detectable in mitotic cells.

### CRISPR/Cas9 knock-in

800 base pair homology arms that span left and right of the start codon of B56α and B56γ were custom synthetized by Biomatik. A NaeI (B56γ)/SwaI (B56α) restriction site was place between the homology arms and used to insert a YFP tag by Gibson assembly. Guides were designed to span the start codon (using http://crispr.mit.edu/). Flp-in HeLa Cas9 cells were generated and transfected with the YFP-homology arm vector and guide in a 1:1 ratio. Cas9 expression was then induced by addition of doxycycline and FACS sorting was performed 2 weeks later to enrich for the YFP-expressing population.

### Live-cell imaging and immunofluorescence

For time-lapse analysis, cells were plated in 24-well plates, transfected and imaged in a heated chamber (37 °C and 5% CO2) using a 10x/0.5 NA on a Zeiss Axiovert 200M Imaging system, controlled by Micro-manager software (open source: https://www.micro-manager.org/). Images were acquired with a Hamamatsu ORCA-ER camera every 4 minutes using 2×2 binning. For immunofluorescence, cells were plated on High Precision 1.5H 12-mm coverslips (Marienfeld). Following the appropriate treatment, cells were pre-extracted with 0.1% Triton X-100 in PEM (100 mM Pipes, pH 6.8, 1 mM MgCl2 and 5 mM EGTA) for 1 minute followed by addition of 4% PFA/PBS for 2 minutes; cells were subsequently fixed with 4% paraformaldehyde in PBS for 10 minutes. Coverslips were washed with PBS and blocked with 3% BSA in PBS + 0.5% Triton X-100 for 30 minutes, incubated with primary antibodies for 16 h at 4 °C, washed with three times with PBS and incubated with secondary antibodies plus DAPI for an additional 2-4 hours at room temperature in the dark. Washed coverslips were then mounted on a glass slide using ProLong antifade reagent (Molecular Probes). All images were acquired on a DeltaVision Core or Elite system equipped with a heated 37°C chamber, with a 100x/1.40 NA U Plan S Apochromat objective using softWoRx software (Applied precision). Images were acquired at 1×1 binning using a CoolSNAP HQ2 camera (Photometrics) and processed using softWorx software and ImageJ (National Institutes of Health). All images displayed are maximum intensity projections of deconvolved stacks. All displayed immunofluorescence images were chosen to most closely represent the mean quantified data.

### Image quantifications

For kinetochore quantification of immunostainings, all images within an experiment were acquired with identical illumination settings and analysed using ImageJ (for experiments in which ectopic proteins were expressed, cells with comparable levels of exogenous protein were selected for analysis). Kinetochore quantification were performed as previously (Saurin et al., 2011). For quantification of B56 localization, a line was drawn through kinetochore pairs lying on the same Z-section (using ImageJ), with the first kinetochore peak at 0.2 µm from the start of the line. An ImageJ macro (created by Kees Straatman, University of Leicester and modified by Balaji Ramalingam, University of Dundee) was used to simultaneously measure the intensities in each channel across the line. The CENP-C channel was used to choose 5 random kinetochore pairs per cell. The signal from the 5 kinetochore pairs was averaged and normalized to the maximum signal in each channel. For chromosome alignment assays, misalignments were score as mild (1 to 2 misaligned chromosomes), intermediate (3 to 5 misaligned chromosomes), and severe (>6 misaligned chromosomes). For mitotic exit assays, time of entry into mitosis (T0, defined by the rounding up of the cell) and the time of anaphase (T1, defined by the separation of the sister chromatids or flattening down of the cell in nocodazole+AZ-3146) were recorded for 50 cells. Data is presented as cumulative percentage of mitotic exit over time.

### Immunoprecipitation and immunoblotting

Flp-in HeLa cells were treated with thymidine and doxycycline for 24h and subsequently released into fresh media supplemented with doxycycline and nocodazole for 16h. Mitotic cells were isolated by mitotic shake off and lysed in lysis buffer (50 mM Tris, pH 7.5, 150 mM NaCl, 0.5% TX-100, 1 mM Na_3_VO_4_, 5 mM ß-glycerophosphate, 25 mM NaF, 10 nM Calyculin A and complete protease inhibitor containing EDTA (Roche)) on ice. The lysate was incubated with GFP-Trap^®^ magnetic beads (from ChromoTek) for 2h at 4 °C on a rotating wheel in wash buffer (same as lysis Buffer, but without TX-100) at a 3:2 ratio of wash buffer:lysate. The beads were washed 3x with wash buffer and the sample was eluted according to the protocol from ChromoTek. Samples were them processed for SDS-Page and immunoblotting using standard protocols.

### Quantification of immunoblots

For quantification of relative immunoprecipitation levels, scanned immunoblots were analyzed using Image Studio Lite (LI-COR Bioscences). A rectangle of the same size was drawn around each band and the intensity within the band (minus the background) was calculated. The immunoprecipitated protein was used as a control, and each band was normalized to it.

### Antibodies

All antibodies were diluted in 3% BSA in PBS. The following primary antibodies were used for immunofluorescence imaging (at the final concentration indicated): mouse α-GFP (clone 4E12/8, a gift from P. Parker; 1:1000), chicken α-GFP (ab13970, Abcam; 1:5000), mouse α- Sgo1 (clone 3C11, H00151648-M01, Abnova; 1:1000), rabbit α-Sgo2 (A301-262A, Bethyl; 1:1000), mouse α-BubR1 (clone 8G1, 05-898, Upstate/Millipore; 1:1000), mouse α-VSV (clone P5D4, V5507, Sigma; 1:1000), rabbit α-Knl1 (ab70537, Abcam; 1:1000), rabbit α-Bub1 (A300-373A, Bethyl; 1:1000), mouse α-FLAG (clone M2, F3165, Sigma, 1:10000) guinea pig α-CENP-C (BT20278, MBL; 1:5000) and rabbit α-pMELT-Knl1 directed against T943 and T1155 of human Knl1(Nijenhuis et al., 2014). Secondary antibodies used were highly-cross absorbed goat α-rabbit, α-mouse, α-guinea pig or α-chicken coupled to Alexa Fluor 488, Alexa Fluor 568, or Alexa Fluor 647 (Life Technologies); all were used at 1:1000.

The following antibodies were used for western blotting (at the final concentration indicated): rabbit α-GFP (custom polyclonal, a gift from G. Kops; 1:5000), mouse α-B56γ (clone A-11, sc-374379, Santa Cruz Biotechnology; 1:1000), mouse α-B56α (clone 23, 610615, BD; 1:1000), mouse α-PPP2CA (clone 1D6, 05-421, Millipore; 1:5000) and rabbit α-PPP2R1A (clone 81G5, #2041, CST; 1:1000), rabbit α-BubR1 (A300-386A, Bethyl; 1:1000), rabbit α-Axin (C76H11, CST; 1:1000), rabbit α-GEF-H1 (155785, Abcam; 1:1000), rabbit α-Kif4a (A301-074A, Bethyl; 1:1000), rabbit α-RepoMan (HPA030049, Sigma; 1:1000), rabbit α-BubR1 pT670 (custom polyclonal, a gift from G. Kops; 1:1000), rabbit α-BubR1 pT680 (ab200061, Abcam; 1:1000). and rabbit α-Actin (A2066, Sigma; 1:5000). Secondary antibodies used were goat α-mouse IgG HRP conjugate (Bio-Rad; 1:2000) and goat α-rabbit IgG HRP conjugate (Bio-Rad; 1:5000).

### Statistical tests

Two-tailed, unpaired *t*-tests with Welch’s correction were performed to compare experimental groups in immunofluorescence quantifications (using Prism 6 software). The comparisons most pertinent for the conclusions are shown in the figures and legends.

